# A comparison of movement-related neuronal activities in cerebellar- and basal ganglia-recipient regions of the macaque thalamus

**DOI:** 10.1101/2025.09.12.675921

**Authors:** Daisuke Kase, Andrew J. Zimnik, Karin M Cox, Thomas Pearce, Robert S. Turner

## Abstract

The ventral lateral (VL) nucleus of the thalamus relays signals from the cerebellum (Cb) and basal ganglia (BG) to primary motor cortex (M1). In primates, glutamatergic Cb efferents from the deep cerebellar nuclei and GABAergic BG efferents from the internal segment of the globus pallidus (GPi) terminate in distinct subregions of VL: the posterior (VLp) and anterior (VLa) divisions, respectively. This anatomical segregation suggests that Cb- and BG-thalamocortical circuits may play distinct roles in motor control, which could be revealed by comparing movement-related activity in VLp and VLa. Here, we recorded single-unit activity from VLp and VLa, identified via electrical stimulation of superior cerebellar peduncle and GPi, during a choice reaction time reaching task. We also recorded from M1, which maintains bidirectional connections with both VLp and VLa. Peri-movement increases in firing began earlier in VLp than in VLa whereas decrease-type responses were more prevalent and prolonged in duration in VLa neurons as compared with VLp and M1. Time-resolved general linear model analysis showed dynamic encoding of task parameters, particularly movement direction, in all three regions. Direction encoding was strongest in M1, moderate in VLp, and weakest in VLa. Direction encoding in VLa also lagged behind that in M1 and VLp. Clustering analysis of direction encoding strength and timing revealed a subpopulation of VLp neurons that encoded direction particularly strongly during the reaction time period. These results highlight a limitation of traditional assumptions that activity characteristics are distributed homogeneously across neural populations and suggest a novel functional organization within VLp neurons.

## 1 Introduction

The learning and production of skilled movement is thought to depend on dynamic interactions between primary motor cortex (M1) (Evarts, 1973; Floyer-Lea and Matthews, 2005; Matsuzaka et al., 2007; Schieber and Rivlis, 2007) and multiple subcortical reentrant loop circuits. Chief among these are circuits through the cerebellum (Cb) (Evarts and Thach, 1969) and basal ganglia (BG) (Alexander and Crutcher, 1990a; Hoover and Strick, 1999). The Cb circuit has been hypothesized to be involved in predictive control, especially of the timing of movement (Ivanusic et al., 2005; Nashef et al., 2019) and correction of sensorimotor error (Fortier et al., 1989; Horne and Butler, 1995; Fu et al., 1997; Kitazawa et al., 1998; Sedaghat-Nejad et al., 2019) whereas the BG circuit may be involved in reward-related selection of action (Mink, 1996; Redgrave et al., 1999), formation of habitual skills (Graybiel, 2008; Ashby et al., 2010), and modulation of motor vigor (Mazzoni et al., 2007; Turner and Desmurget, 2010; Yttri and Dudman, 2016). Both subcortical circuits relay signals to primary motor cortex (M1) by way of neurons in the ventral lateral (VL) nucleus of the thalamus. In primates, Cb efferents from the deep cerebellar nuclei (DCN) and BG efferents from the internal segment of the globus pallidus (GPi) terminate in distinct posterior and anterior parts of VL (VLp and VLa, respectively) (Nauta and Mehler, 1966; DeVito and Anderson, 1982; Asanuma et al., 1983; Ilinsky and Kultas-Ilinsky, 1987) such that, in primates, convergence of inputs from the DCN and GPi onto individual VL neurons is rare (Yamamoto et al., 1983; Anderson and Turner, 1991; Schwab et al., 2020), but see (Sakai et al., 1996; Koster and Sherman, 2024; Roth et al., 2024). This anatomical separation of the DCN- and GPi-recipient thalamic neurons suggests that the distinct roles in motor control hypothesized for Cb- and BG-thalamocortical circuits may be evident through a comparison of the movement-related activities of neurons in VLp and VLa.

The DCN- and GPi-thalamic circuits share many similarities. The synapses of both onto their respective VL thalamo-cortical neurons have morphologic features characteristic of giant “driver-type” synapses (Sherman and Guillery, 1996; Kultas-Ilinsky et al., 2003; Rovo et al., 2012). Electrical stimulation applied to either of the two projections evokes strong responses in recipient VL neurons, effects interpreted as evidence of a driver-type synaptic function (Uno et al., 1970; Nambu et al., 1991; Ando et al., 1995; Kim et al., 2017); however, see (Kase et al., 2015). In addition, projection neurons in both DCN and GPi have high firing rates at rest (both typically 50 spikes/sec or higher in primates)(Thach, 1970; DeLong, 1972; Mink and Thach, 1991). One critical difference, however, is that DCN→VLp synapses are glutamatergic whereas GPi→VLa synapses are GABAergic (Uno et al., 1970; Uno et al., 1978; Penney and Young, 1981; Rovo et al., 2012).

A default perspective would be that the thalamocortical neurons of VLp and VLa simply relay to cortex the input signals they receive from the Cb and BG, respectively. That view predicts that single-unit activity in VLp should reflect the combined effects of high tonic excitation received from DCN projection neurons and modulations away from that high baseline reflecting the contributions of Cb output to motor cortical function. In contrast, activity in VLa should show the combined effects of high frequency resting inhibition from GPi and phasic modulations in firing rate that reflect BG contributions to motor cortex. It is quite surprising then that the few studies to have investigated this question found only minor differences between the two nuclei with respect to single-unit baseline firing rates or patterns (Pessiglione et al., 2005), but see (Vitek et al., 1994), and the general sign and timing of movement-related modulations in activity (Anderson and Turner, 1991; van Donkelaar et al., 1999). In the rodent motor thalamus, segregation of Cb- and BG-recipient neurons into distinct anatomic regions is less obvious or complete than in primates (Bosch-Bouju et al., 2013; Phillips et al., 2019; Koster and Sherman, 2024; Roth et al., 2024) although see (Kuramoto et al., 2011). As a result, few neurophysiologic studies of motor thalamus in the rodent have distinguished Cb- and BG-recipient populations (Bosch-Bouju et al., 2014; Guo et al., 2017; Gaidica et al., 2018; Sauerbrei et al., 2020; Inagaki et al., 2022).

The conventional view of motor thalamus as a relay for cortex-bound subcortical inputs sidesteps the abundant anatomic and physiological evidence that thalamic nuclei are also influenced strongly by inputs from cortex mediated via mono-synaptic excitatory inputs (Rouiller et al., 1998; Galvan et al., 2016), via disynaptic inhibitory inputs through the reticular nucleus of the thalamus (Ilinsky et al., 1999; Jones, 2007) and, in primates, via GABAergic intra-thalamic interneurons (Ilinsky and Kultas-Ilinsky, 1990; Arcelli et al., 1997; Galvan et al., 2016). The anatomy and physiology of these circuits suggest that VLp and VLa are likely to integrate their respective ascending inputs with descending signals received from cortex. While the segregation in the primate thalamus of inputs from the Cb and BG suggest distinct computational roles for VLp and VLa, the extent to which these pathways are engaged in different dimensions of behavior has seldom been investigated and, as mentioned above, the extant studies described surprisingly minor differences between VLp and VLa (Nambu et al., 1988; Anderson and Turner, 1991; van Donkelaar et al., 1999).

To clarify the roles in motor control of Cb- and BG-thalamocortical circuits, we compared the activity of neurons in VLp and VLa during performance of a choice reaction time reaching task. Following existing evidence (Nambu et al., 1988; Anderson and Turner, 1991; van Donkelaar et al., 1999), we predicted that general characteristics, such as baseline firing rates, and the timing and sign of movement-related modulations (i.e. increases or decreases in firing rate), would not differ significantly between VLp and VLa. We suspected, however, that a more detailed examination of the relationships between peri-movement activity and measures of task performance would reveal differences between VLp and VLa populations consistent with the computational roles proposed for their respective Cb and BG inputs. To address these hypotheses, we studied the single-unit activity of electrophysiologically-identified neurons in the VLp and VLa of two non-human primates during performance of a simple well-learned reaching task. We evaluated the dynamic encoding of individual parameters of task performance using a time-resolved general linear model (GLM) analysis. For added context, we compared those results against single-unit data collected from the proximal arm region of M1. In most respects, our results confirm previous reports of overall similarity in firing properties between the two nuclei and extend those findings to general features of the time-resolved encoding of movement parameters. Those results suggest that common inputs to both thalamic regions from motor cortical areas dominate many aspects of the activity of both VLp and VLa, at least in the context of a well-learned highly stereotyped reaching behavior. The regression analysis did reveal a subpopulation of VLp neurons that encoded movement direction particularly strongly during the reaction time interval. In that VLp population, direction-related semi-partial R^2^ reached higher values than those seen in VLa and M1 neurons. In addition, we did find that encoding of direction in VLa lagged behind what was observed in VLp and M1. These results provide clues to the different roles in motor control played by cerebello-thalamic and basal ganglia-thalamic circuits.

## 2 Materials and Methods

### 2.1 Animals and behavioral task

Two monkeys (Macaca mulatta; G, female 7.1kg; I, female 7.5kg) were used in this study. All aspects of animal care were in accord with the National Institutes of Health Guide for the Care and Use of Laboratory Animals, the PHS Policy on the Humane Care and Use of Laboratory Animals, and the American Physiological Society’s Guiding Principles in the Care and Use of Animals. All procedures were approved by the Institutional Animal Care and Use Committee. The animals performed a choice reaction time reaching task (Fig. 1A) that has been described in detail previously (Franco and Turner, 2012; Zimnik et al., 2015) (DOI: 10.5281/zenodo.11397910). In brief, the animal faced a vertical response panel that contained two target LEDs, positioned 7 cm to the left and right of midline. To begin each trial, the animal’s left hand rested at a “home-position”, a platform located lateral to the left hip at waist height and equipped with a proximity sensor. The animal was trained to hold the home-position (2–10s, uniform random distribution) until illumination of one of the target LEDs, selected in pseudo-random order trial-to-trial, served as both a “Go” signal and a directional instruction. To successfully complete the trial, the animal was required to move its hand from the home-position to the target within 1s of appearance of the Go cue and hold the hand at the target position for 0.6-1.2s (random uniform distribution). After the target hold condition was met, a drop of pureed food was delivered via a sipper tube and computer-controlled peristaltic pump. Following reward delivery, the animal was allowed, with no time constraints, to return its hand to the home-position and thereby initiate the next trial. A trial was marked as an error and reward was not delivered if the animal failed to complete the hold period, reached to the incorrect target, or if the 1 sec response window was exceeded. The presence of the hand at the home-position and at targets was detected by infrared proximity sensors (Takex, GS20N).

**Figure 1.**
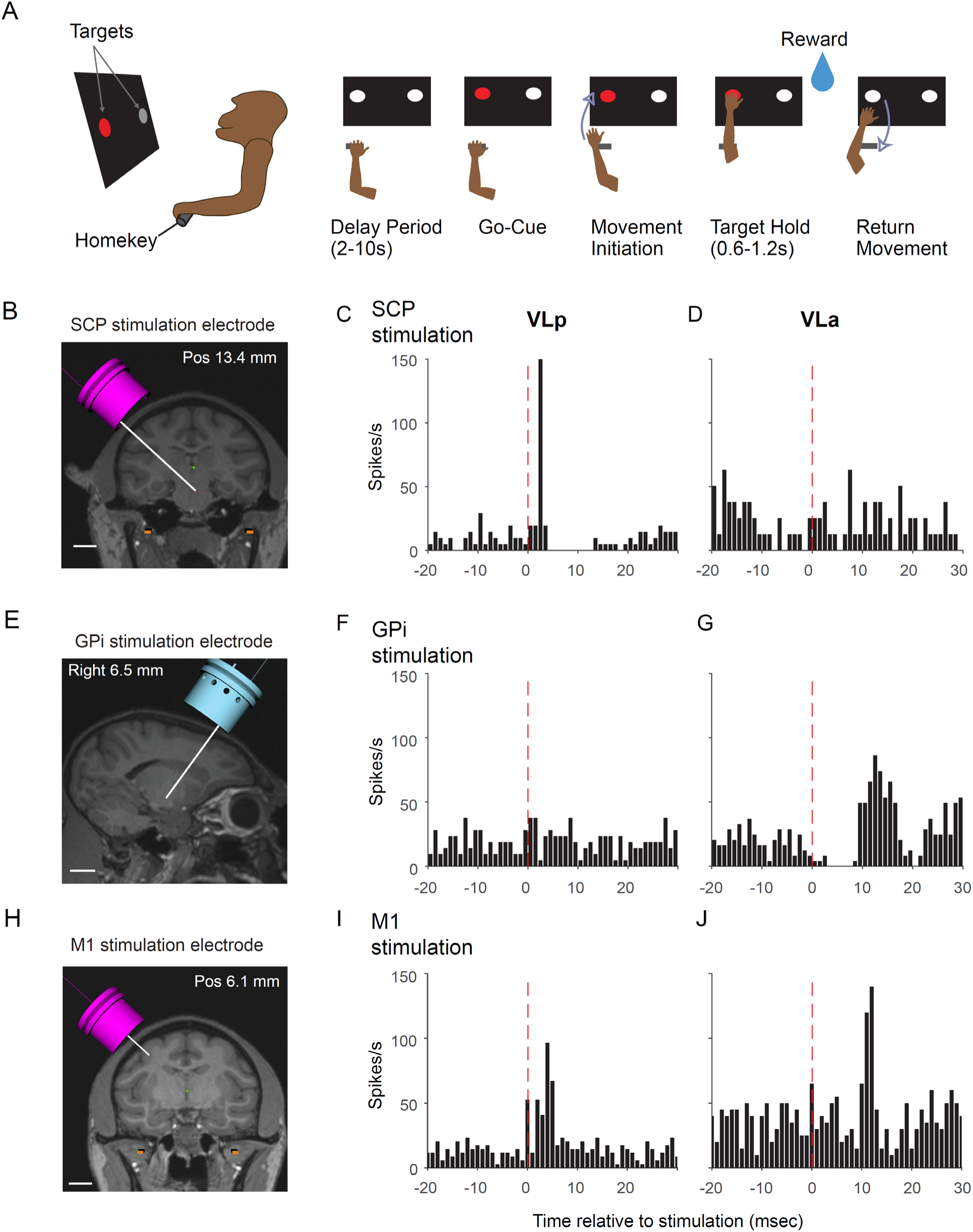
**(A)** Schematic of the task. **(B), (E) and (H)** Schematic illustration of the stimulation electrode tracks targeting the superior cerebellar peduncle (SCP) (B), GPi (E), and M1 (H). Scale bar: 10 mm. **(C-D, F-G, and I-J)** Exemplar peri-stimulus time histograms of neuronal spike activity averaged around the times of stimulation (time 0) for a single unit located in VLp (C, F, and I) and a unit located in VLa (D, G, and J). The top panels (C-D) show responses to SCP stimulation, the middle panels (F-G) show responses to GPi stimulation, and the bottom panels (I-J) show responses to M1 stimulation.

### 2.2 Surgery

General surgical procedures have been described previously (Desmurget and Turner, 2008; Zimnik et al., 2015). The chamber implantation surgery was performed under sterile conditions with ketamine induction followed by Isoflurane anesthesia. Vital signs (pulse rate, blood pressure, respiration, end-tidal pCO_2_, and EKG) were monitored continuously to ensure proper anesthesia. A cylindrical titanium recording chamber was affixed to the skull at stereotaxic coordinates to allow access to the right globus pallidus and ventrolateral thalamus via a parasagittal approach. A second chamber was positioned over the right hemisphere in the coronal plane to allow access to the arm area of primary motor cortex (M1) and, in the midbrain, the decussation of the superior cerebellar peduncle (SCP). The chambers and head stabilization devices were fastened to the skull via bone screws and methyl methacrylate polymer.

Prophylactic antibiotics and analgesics were administered post-surgically. We have uploaded a more detailed description of procedures to protocols.io (implantation of head fixation and recording chamber https://dx.doi.org/10.17504/protocols.io.5jyl8p9yrg2w/v1 and craniotomy https://dx.doi.org/10.17504/protocols.io.n92ldmz48l5b/v1).

### 2.3 Localization of stimulation sites and implantation of indwelling macroelectrodes

To allow electrical stimulation-based identification of the DCN-recipient VLp and the GPi-recipient VLa (Anderson and Turner, 1991), we implanted stimulation electrodes chronically in the SCP at its decussation and in the arm-related region of the primary motor cortex (Figs. 1B and 1H). The anatomic locations of sites for implantation were estimated initially from structural MRI scans (Siemens 3T Allegra Scanner, voxel size of 0.6mm) using an interactive 3D software system (Cicerone) to visualize MRI images and predict trajectories for microelectrode penetrations (Miocinovic et al., 2007). Precise coordinates for each target were then determined using microelectrode mapping methods. And then, custom-built stimulating electrodes were implanted at those sites using methods described previously (Turner and DeLong, 2000). The custom macroelectrodes consisted of two Teflon-insulated Pt-Ir microwires (50µm) glued inside a short stainless-steel cannula with ∼0.5mm separation between the distal ends of the microwires. Insulation was stripped from ∼0.2mm of the distal ends of the microwire to achieve an impedance of ∼10kΩ. Electrode assemblies for SCP and M1 targets were implanted transdurally via the coronal chamber using a protective guide cannula and stylus mounted in the microdrive. In the months following implantation, the location and integrity of macroelectrodes were monitored by comparing electrode impedances and the current thresholds for activation and the specific muscle contractions evoked by stimulation through each electrode.

### 2.4 Localization of subcortical target regions for recording and stimulation

Many of the target localization methods have been described previously (Schwab et al., 2020). In brief, the chamber coordinates for candidate regions were estimated initially from structural MRIs as described above. Single-unit microelectrode recording was then performed in combination with electrical stimulation (single biphasic pulses <200μA, 0.2 ms-duration at 2Hz max.; Model 2100, A-M Systems) and proprioceptive exams performed by the experimenter.

#### Globus Pallidus interna

To locate the boundaries of the GPi-recipient VLa nucleus of thalamus, we first needed to identify an appropriate location in GPi for electrical stimulation. The target location in GPi was identified by the presence of typical high firing rate single-units, many of which responded briskly to a) proprioceptive stimulation of the forelimb (Turner and Anderson, 1997) and b) electrical stimulation in the arm region of primary motor cortex (Yoshida et al., 1993). We placed a stimulating electrode acutely at this GPi location during the subsequent mapping of VLa boundaries (Fig. 1E).

#### VL thalamus

The general boundaries of VL thalamus were identified using stereotactic planning under MR guidance and then electrophysiological confirmation as described in (Anderson and Turner, 1991; Vitek et al., 1994; Buford et al., 1996). Subregions of the VL that were arm-related and likely to be connected with M1 were defined by a) the presence of unit activity typical of thalamus that was modulated by voluntary or passive movements of the animal’s proximal arm and b) observation, in a subset of those units, of short latency responses to electrical stimulation of one or more of the macroelectrodes implanted in M1 (Fig. 1I and J).

#### VLp thalamus

Within the VL nucleus, as defined above, VLp was identified by its position (Ilinsky and Kultas-Ilinsky, 1987) and the presence of neuronal discharge that: a) responded at short latency to SCP stimulation (Fig. 1C), and b) did not respond to GPi stimulation (Fig. 1F).

#### VLa thalamus

VLa was identified by its characteristic position in the anterior-lateral part of VL and the presence of single-unit discharge that: a) responded to GPi stimulation with a short latency pause in firing (Fig. 1G)(Nambu et al., 1988; Anderson and Turner, 1991), often followed by a rebound increase in firing probability, and b) showed no response to SCP stimulation (Fig. 1D).

#### Thalamic microstimulation

Also, we tested the electrical excitability of VL recording sites in accord with previous reports that low current microstimulation evoked muscle contractions only from locations within VLp and not from VLa (Vitek et al., 1994; Buford et al., 1996). We delivered short trains of high frequency electrical stimulation (glass-insulated tungsten microelectrodes; 10 biphasic pulses <200μA, 0.2 ms-duration at 300Hz; Model 2100, A-M Systems) at sites classified as VLa and VLp according to the criteria above. Consistent with previous reports, stimulation at many locations classified as VLp evoked movement at 40-60 μA thresholds (Vitek et al., 1994; Buford et al., 1996), while stimulation at locations classified as VLa rarely evoked movement. On occasion, however, we evoked muscle contractions from VLa sites that were located near the border with VLp, making microstimulation an imprecise marker of the boundary between nuclei. Therefore, microstimulation findings were not used as a primary criterion for identification of the VLa/VLp border.

#### Primary motor cortex (M1)

Single-unit recordings were obtained from the M1 of one animal (monkey I). M1 sites for single-unit recording were localized by identification of the principal arm territory along the anterior bank and gyrus of the pre-central sulcus. As reported extensively, typical neuronal discharge responded to active and/or passive movement of the arm and microstimulation at low current (<60 μA, 10 biphasic pulses at 300 Hz) evoked contraction of forelimb muscles (Asanuma and Rosen, 1972). Single-unit data from the M1 of the other animal were not available due to technical problems.

### 2.5 Recording protocol

The extracellular spiking activity of neurons in VLp, VLa and M1 was sampled using multiple glass-insulated tungsten microelectrodes (0.5–1.5MΩ, Alpha Omega Co.) or 16-contact linear probes (0.5–1.0MΩ, V-probe, Plexon Inc.). Probes were introduced transdurally protected within custom-sharpened cannulae (Vita Needle Company, Inc.). Voltage signals from each electrode contact were amplified, band-pass filtered (4×, 2Hz–7.5kHz), and digitized at 24 kHz (16-bit resolution; Tucker Davis Technologies). To enhance the mechanical stability of recordings, the 16-contact linear probes were lowered initially to a depth 1mm below the target depth, then withdrawn and left at the target depth for 30min before initiating data collection. When stable single-unit isolation was available from one or more single-units in either VLp or VLa, as determined by online spike sorting, neuronal data and behavioral events were collected while the animal performed the behavioral task. On protocols.io, we have posted protocols relevant to electrophysiological recording (https://dx.doi.org/10.17504/protocols.io.bp2l6xx91lqe/v1) and recording chamber maintenance (https://dx.doi.org/10.17504/protocols.io.5jyl8ppm9g2w/v1).

### 2.6 Offline analysis

#### 2.6.1 Behavior

Task performance was screened and error trials and outliers in task performance were excluded from further analyses. Error trials included reaches to the incorrect target and failure to reach the target within the allowed 1s interval after the go cue. Two behavioral metrics, reaction time and movement duration, were used for outlier detection. Reaction time (RT) was defined as the interval between the go cue illumination and the lift of the hand off the home position as detected by the infrared sensor, termed movement onset. Movement duration (MD) was defined as the interval between movement onset and target capture, which was detected by an infrared sensor at the target. Trials were classified as outliers if either behavioral metric exceeded a threshold of 6 median absolute deviations from the median (Matlab: ISOUTLIER). For trials that met the above inclusion criteria, average movement speed was computed by dividing the distance from the home position to the target (left: 35cm; right: 42cm) by the MD.

#### 2.6.2 Spike sorting and detection of peri-movement discharge

The stored wide-band neuronal signal was high-pass filtered (Fpass: 300Hz, Matlab FIRPM), manually thresholded, and candidate action potentials were sorted into clusters in principal components space (Off-line Sorter, Plexon Inc.). Clusters were accepted as well-isolated single-units only if the unit’s action potentials were of a consistent shape and could be separated reliably from the waveforms of other neurons as well as from background noise throughout the period of recording. Single-units from VLp, VLa and M1 were included for further analysis if they exceeded minimum firing rate thresholds (0.9Hz for VLp and VLa, 0.1Hz for M1, averaged over the entire recording session). A minimum of 20 valid behavioral trials (minimum of 10 in each direction) was required to include a unit in task-based analyses.

Measures of baseline activity were calculated from neural activity sampled during start position hold period before the presentation of the go cue. Measurements of the spike waveform were calculated from spike data that were up-sampled to 244 kHz using linear interpolation (Matlab INTERP1) and then averaged. Action potential width was calculated as the full width at half-maximum of the first trough. Burst-like events were detected using the Poisson surprise method as described previously ((Legendy and Salcman, 1985); initial filtering threshold factor: 1.5, minimum spike number of the burst: 3, surprise cutoff: 5).

We tested for peri-movement changes in single-unit spike rate using a standard rate change detection method (Alexander and Crutcher, 1990b; Zimnik et al., 2015). First, spike density functions (SDFs) were constructed by convolving each unit’s spike time stamps (1 kHz resolution) with a Gaussian kernel (σ = 25 ms). SDFs were then aligned to the time of movement onset for each trial and across-trial mean SDFs were constructed separately for reaches to each target. The baseline activity for each unit was calculated as the mean and standard deviation of the SDF over a 700 ms window ending at the median time of target LED onset, correcting for linear trends in the SDF over this window. This baseline activity was then used to detect movement-related changes in the SDF during a peri-movement period that extended from the median time of go cue onset (i.e., illumination of the target LED) until the median time of target capture, thereby encompassing both reaction time and movement duration intervals. We defined a movement-related change in activity as a statistically-significant elevation or depression in the SDF from baseline that lasted at least 60 ms (e.g., Fig.2 A, solid vertical lines; t-test, one-sample vs. baseline; omnibus p < 0.001 after Bonferroni correction for multiple comparisons). All movement-related changes in firing rate were classified as increases or decreases in discharge rate and the time of onset of that change was defined as the first time point in the SDF that crossed the threshold for significance. Note that this approach enabled detection of biphasic changes (e.g., an increase followed by a decrease), but further analysis was based only on the earliest-occurring peri-movement change in firing.

**Figure 2.**
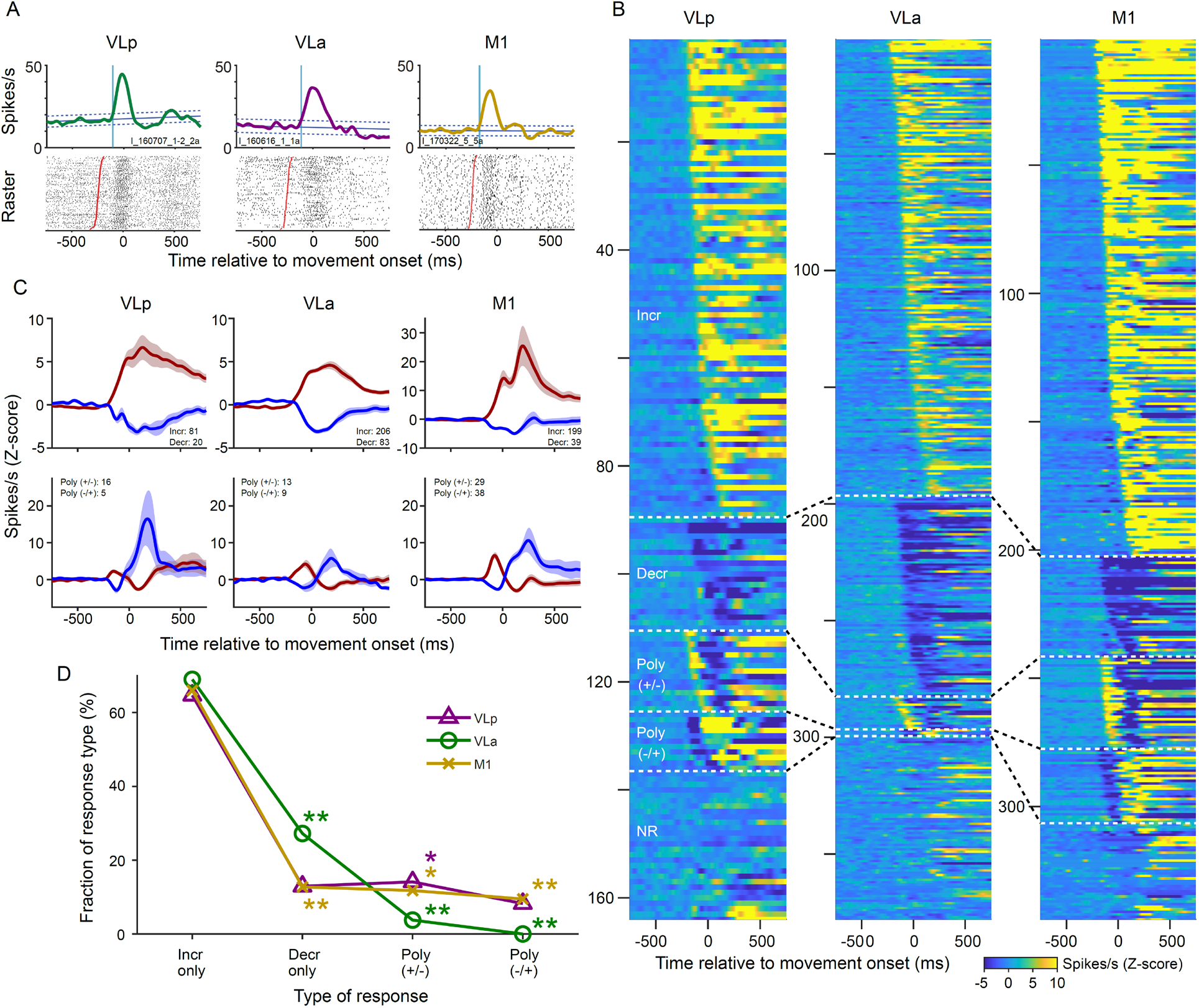
**(A)** Peri-movement activity of exemplar single units sampled from VLp (left), VLa (middle), and M1 (right). Across-trial mean spike density functions (SDFs; top) and raster plots (bottom) are aligned to movement onset. *Sloped blue lines:* the unit’s mean baseline trendline ± confidence interval. *Vertical cyan line*: the estimated onset of the mean response. In rasters, single-trial spike trains are sorted by reaction time (shortest at top). Red circles indicate go-cue onset for each trial. **(B)** Peri-movement SDFs of all recorded single units, sorted by response type and onset latency (earlier responses at the top within each category) in VLp (left), VLa (middle), and M1 (right). SDFs were z-scored relative to mean baseline activity prior to the go-cue onset. **(C)** Population averages of z-scored SDFs. Top: monophasic responses (red: increases only; blue: decreases only). Bottom: polyphasic responses (red: increase–decrease [+/-]; blue: decrease–increase [-/+]). Data shown for VLp (left), VLa (middle), and M1 (right). *Shaded areas*: ±SEM. **(D)** Proportions of peri-movement response types in VLp (magenta), VLa (green), and M1 (cyan). The proportion of decrease-only responses in VLp and M1 was significantly smaller than the proportion estimated from all responses, whereas the proportion of polyphasic responses in VLp and M1 was significantly larger than expected (*p<0.05, **p<0.01; standardized residual analysis). Magenta and cyan asterisks denote significant effects in VLp and M1, respectively.

#### 2.6.3 Time-resolved general linear model (GLM) regression

We used general linear model regression to determine if single-unit firing rate encoding of measures of task performance differed between VLp and VLa neurons. We used the Matlab command ‘lassoglm’ with a normal regression model, 10-fold cross validation (to estimate the reliability of results) and an ‘alpha’ set to 0.5, thereby implementing elastic-net regularization of potential covariation between regressors. The regressors included trial-by-trial values for parameters controlled by the task (target direction and duration of the start position hold period before go-cue presentation), measures of how the animal performed the task (reaction time and mean movement speed), and the time elapsed since the beginning of the recording session.

Target direction was coded as +1 (right, i.e., contralateral to the working arm) and -1 (left), and the remaining regressors were z-scored to facilitate the comparison of results between regressors. We assessed collinearity using Belsley’s collinearity diagnostics with the Matlab command ‘collintest’ (Belsley, 1980). A condition index threshold of 10 was used to indicate acceptable level of low collinearity (Belsley, 1991). To ensure that the resulting model was sparse, the regularization penalty value lambda was set to the greatest value within one standard deviation of the lambda that minimized cross-validation error. The neural response function to be predicted was a time series of one unit’s spike counts across peri-movement time for each trial. More specifically, we calculated spike counts within a 200 ms wide window at 1200 time points starting centered at 700 ms before movement onset and swept in 1ms steps to 500 ms after movement onset. An independent lassoglm was run on the trial-by-trial spike counts at each 1 ms time step of the response function. Coefficients of semi-partial determination (semi-partial R^2^ values, pR^2^) for each regressor were defined as the proportion of across-trials variation in spike counts explained by the full regression (sum of squared errors) that could not be explained by a reduced model that excluded that regressor.

Then, for each single-unit, we estimated the timing of peri-movement changes in the encoding of individual regressors. We tested for significant increases in a regressor’s pR^2^ during a peri-movement test period that extended from the latest time across trials of go-cue presentation relative to movement onset (i.e., beginning at the shortest RT prior to movement onset) to the earliest end of reach (i.e., ending at the shortest MD following movement onset). If the maximum semi-partial R^2^ within the test period reached a value greater than 0.01 then it was classified as a significant increase in semi-partial R^2^. The time of onset of that increase was defined as the time immediately prior to the maximum when the semi-partial R^2^ crossed a 5% of maximum threshold.

#### 2.6.4 Cluster analysis with Gaussian mixture model

We tested the GLM results for sub-groups (clusters) using Gaussian mixture models (Matlab function: fitgmdist). Those models used full covariance matrices, 10000 iterations, 0.01 of the regularization value, and the number of clusters tested for ranged from one through five. The best-fit number of clusters was identified as the number associated with a minimum in the Bayes Information Criterion (BIC) calculated in the Matlab function (fitgmdist).

## 3 Results

### 3.1 Basic approach and task performance

We studied the single-unit activity of neurons sampled from VLp and VLa of two macaque monkeys and from M1 of one macaque monkey during skilled performance of a choice reaction time reaching task for food reward (Franco and Turner, 2012; Zimnik et al., 2015; Schwab et al., 2020; Cox et al., 2024) (Fig. 1A). Animals performed the reaching task in a highly stereotyped fashion. Reaction times (248.1 ± 33.9 ms and 237.8 ± 42.7 ms for monkey G and I respectively; mean ± SD) and movement durations (249.8 ± 36.0 ms and 199.6 ± 32.8 ms for monkey G and I respectively; mean ± SD) were short and showed a low level of variability. Animals made very few errors in task performance (success rates of 97.5% and 96.3% for monkey G and I, respectively), and the rare error trials were excluded from further analysis.

### 3.2 Identification of VLp and VLa

One or two microelectrodes, or one 16-contact linear probe, were positioned acutely in the arm-related regions of one or the other thalamic nucleus or, occasionally, both nuclei. We used a combination of electrophysiological techniques to classify the recording site as described previously (see Methods, (Schwab et al., 2020)). In brief, VLp regions were identified by the presence of neurons that were excited at short-latency in response to electrical stimulation of the SCP (Fig. 1C) and were not inhibited affected by electrical stimulation of GPi (Fig. 1F). In contrast, VLa regions were identified by the presence of neurons that were inhibited at short-latency following stimulation of GPi (Fig. 1G) and were not excited by stimulation of SCP (Fig. 1D). In both areas, we observed short-latency excitation following electrical stimulation of M1 (Figs. 1I and J). For most recording tracks, the locations of borders between the VLp and VLa regions were determined during preliminary mapping penetrations performed prior to data collection (supplemental Fig. S1). The sub-regions of VLp and VLa involved in arm movement were identified by the presence of short-latency spiking responses to stimulation of arm-related areas of M1. Based on their physiologic responses during mapping, neurons assigned to VLp and VLa categories were found in separate regions of thalamus. Anatomic intermixing of neurons categorized as VLp and VLa was exceedingly rare, consistent with previous reports (Nauta and Mehler, 1966; DeVito and Anderson, 1982; Ilinsky and Kultas-Ilinsky, 1987; Schwab et al., 2020). In rare cases, a neuron encountered close to the electrophysiologically-defined boundary of VLp and VLa territories was both excited by SCP stimulation and inhibited by GPi stimulation. Similar observations have been described previously (Nambu et al., 1988). To avoid potential ambiguities associated with studying such “boundary neurons,” we restricted all subsequent analyses to cells located at least 50um away from the electrophysiologically defined border.

### 3.3 Neuronal recordings and activity at rest

A total of 83 single-units were sampled from the VLp, and 193 single-units were sampled from VLa according to the criteria defined above (see *Methods* for more detailed descriptions). 172 neurons were sampled from the proximal arm-related region of M1 of one animal and are included here for comparison. Units were studied over the course of 132.8 ± 77.3 trials of the behavioral task (mean ± SD) with a mean of 66 trials for each of the two movement directions.

Contrary to expectations based on the distinct synaptic inputs received by VLp and VLa neurons, past studies of the baseline activities of those two populations found negligible differences (Anderson and Turner, 1991; Pessiglione et al., 2005). To revisit that question, we quantified neuronal firing properties across each trial’s start position hold period, which amounted to multisecond-long periods of attentive immobility. Consistent with some (Anderson and Turner, 1991; Pessiglione et al., 2005) but not all (Vitek et al., 1994) previous reports, we found that mean firing rates during the hold period were statistically indistinguishable between VLp and VLa neurons (Wilcoxon rank test; p=0.29; Table 1). Measures of firing rate variability such as the coefficient of variation of inter-spike intervals (CV), proportion of spikes in the burst activity, and the normalized intra-burst firing rate also did not differ significantly between VLp and VLa neurons (Wilcoxon rank test; p=0.38, 0.48, and 0.14 respectively; Table 1). In addition, two measures of action potential duration (half width of the first trough and trough to peak time) did not differ between nuclei (Wilcoxon rank test; p=0.29 and 0.58, respectively; Table 1), again contrary to expectation (Phillips et al., 2019). The ratio of trough magnitudes to peak magnitudes did not differ between nuclei (Wilcoxon rank test; p=0.94).

**Table 1.**
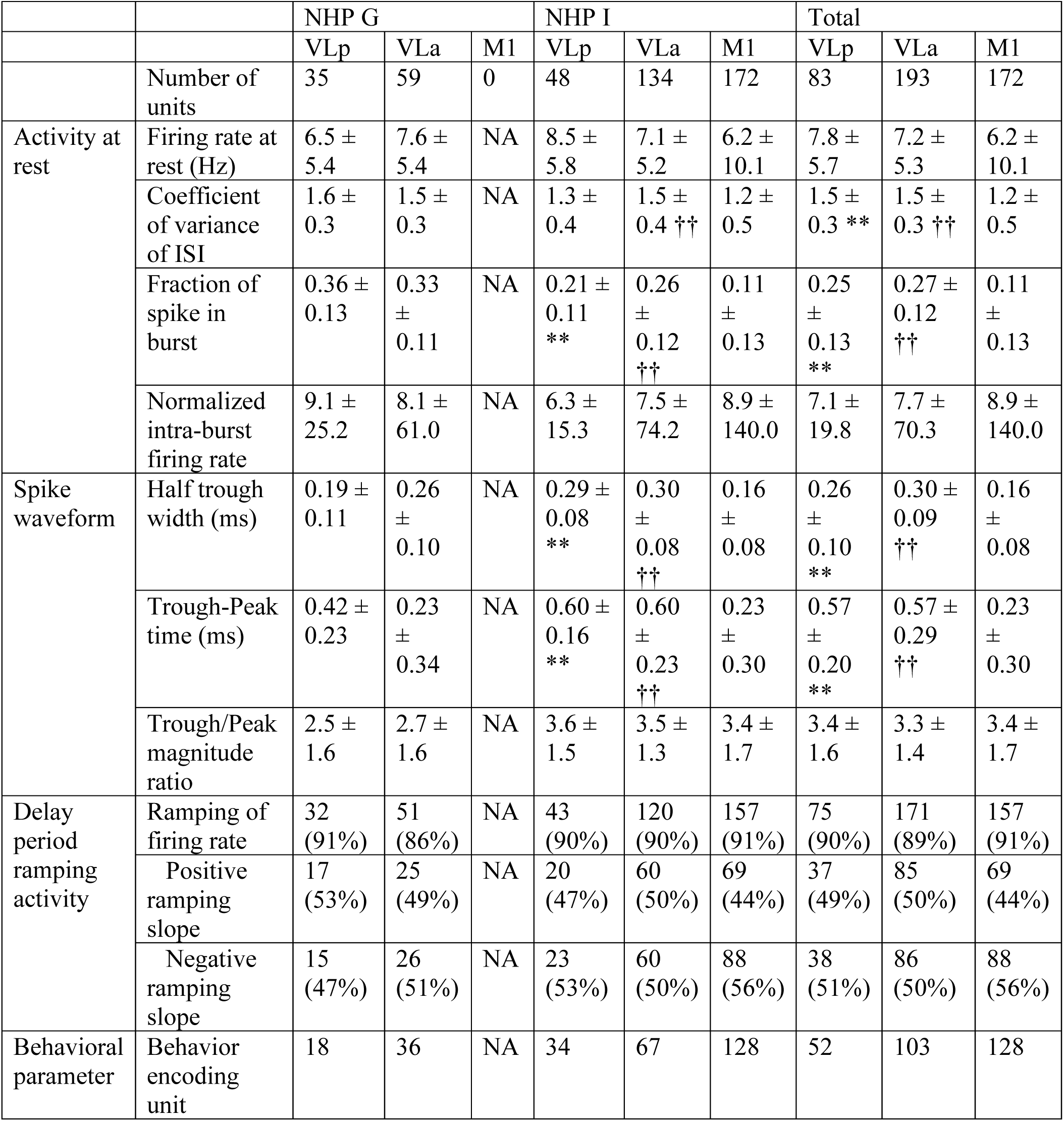
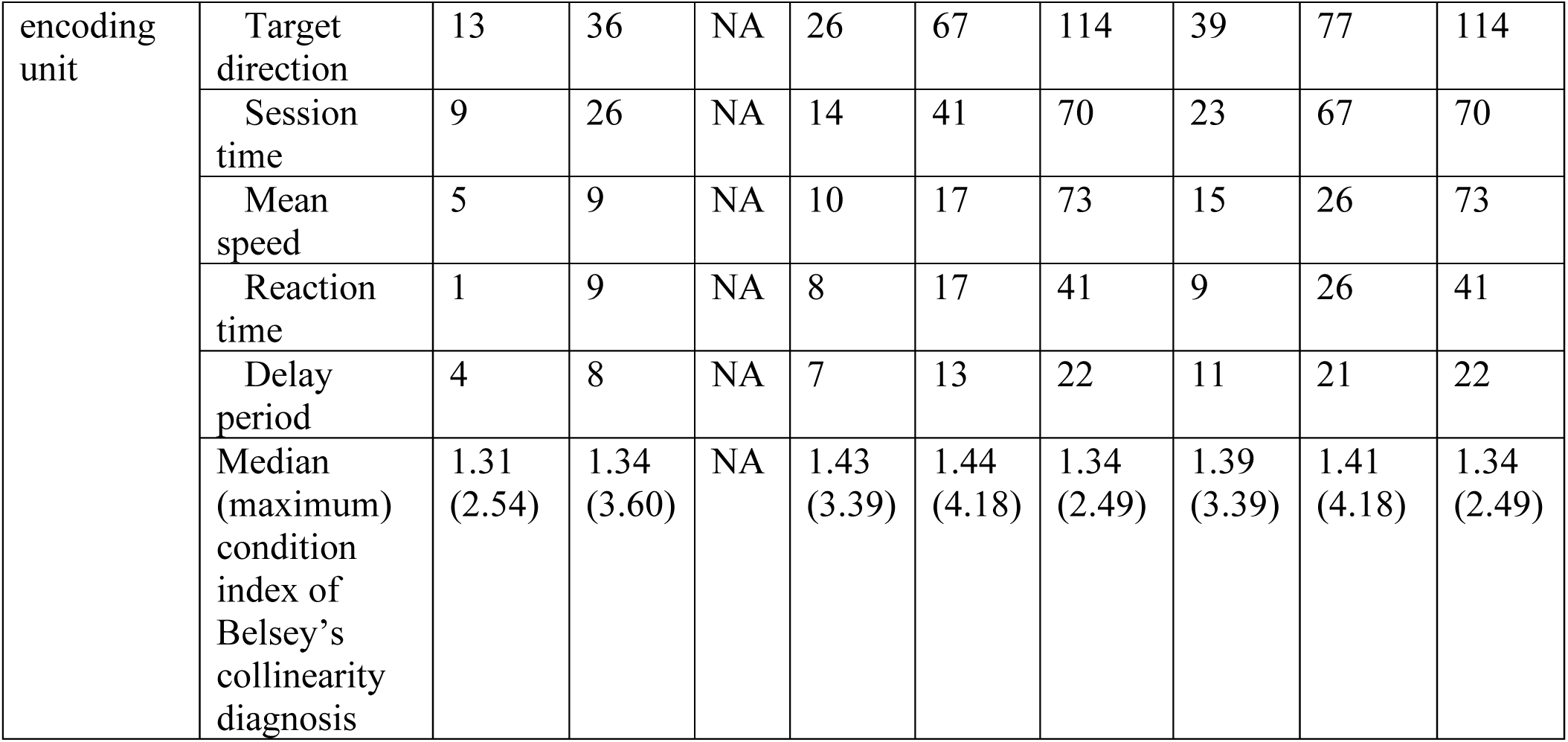
Metrics of activity at rest, spike waveform, delay period ramping activity, and fractions of behavioral parameters encoding unit (median ± MAD, ** p<0.01 between VLp and M1, †† p<0.01 between VLa and M1, Wilcoxon rank test)

Relative to the activity of M1 neurons, neither thalamic nucleus differed significantly in resting firing rates (Wilcoxon rank test; p=0.42 and 0.80 for VLp and VLa respectively). Thalamic neurons, however, were more likely than M1 neurons to fire action potentials in irregular and bursty firing patterns, as evidenced by elevations of the CV of inter-spike intervals and the fraction of spikes in the burst (Wilcoxon rank test; p<0.01; Table 1). Finally, consistent with previous work (Steriade and Llinas, 1988; Sherman and Guillery, 2002), the action potentials of thalamic neurons were prolonged in duration as compared with M1 neurons (half width of the first trough and trough-to-peak times; Wilcoxon rank test; p<0.01). Considering data from NHP I separately (i.e., the animal M1 neurons were recorded from) the trough-to-peak magnitude ratio was larger in thalamic neurons than in M1 (Wilcoxon rank test; p<0.01 and p<0.05 for VLp and VLa respectively; Table 1).

In summary, the baseline activities of neurons in VLp and VLa were indistinguishable from each other in nearly every respect, despite the markedly different subcortical inputs the two thalamic nuclei receive. The baseline activity found in both thalamic nuclei differed from that of M1 across multiple measures in ways that were consistent with previously described generic differences between cortical and thalamic extracellular activity (Steriade and Llinas, 1988; Sherman and Guillery, 2002).

### 3.4 Peri-movement responses in VLp and VLa differ in prevalence, latency and duration depending the sign of the response

Next, we compared the general features of peri-movement modulations in firing of neurons in VLp and VLa. We predicted that the sign and timing of movement-related modulations in activity would not differ substantially between nuclei, consistent with previous reports (Anderson and Turner, 1991; van Donkelaar et al., 1999), but contrary to the common assumption that task-related activity in VLp and VLa is determined primarily by their respective ascending glutamatergic inputs from DCN and GABAergic inputs from GPi. Any differences in response prevalence, form or timing may provide clues into the influences in VL thalamus of those distinct subcortical inputs.

To that end, we tested mean spike density functions (SDFs; see *Methods*) for peri-movement changes in firing relative to a unit’s baseline activity (i.e., linear trend ± SD of activity prior to go-cue presentation, *blue slanted lines* in Fig. 2A). Significant peri-movement changes were detected in large fractions of all neural populations (89%, 89% and 96% in VLp, VLa and M1, respectively (chi-square test; *χ^2^*(2, n=448) = 0.74, p=0.69). The shape and timing of those responses differed widely between neurons and occasionally between directions of movement. We observed both monophasic and poly-phasic responses in all areas. Monophasic responses were composed of a single increase or decrease in firing (Fig. 2 B-D; *Incr* and *Decr*, respectively). Poly-phasic responses were classified according to the sign of the initial response (i.e., an initial increase or decrease in firing rate; Fig. 2 B-D; *Poly (+/-)* and *Poly (-/+)*, respectively). All detected responses are shown in Fig. 2B sorted according to brain area, response type, and latency of response onset relative to the time of movement initiation (time 0). Population averages of these response types for each brain area are shown in Fig. 2C. The rate of occurrence of the four general response types differed between the three brain areas (chi-square test; *χ^2^*(3, n=604) = 32.78, p<0.01; Fig. 2D). Mono-phasic decreases were more common in VLa than in VLp or M1 whereas the poly-phasic decrease-increase response type was less common in VLa than in VLp or M1 (p<0.01, post-hoc adjusted standardized residual analysis, (Haberman, 1973)). In addition, the poly-phasic increase-decrease type of response was less common in VLa than in M1 (p<0.05, adjusted standardized residual analysis).

We measured the properties of individual monophasic responses and then compared them between VLp, VLa and M1 populations (Table 2). (The small numbers of polyphasic responses detected, particularly in VLa, prevented a similar comparison of those response types.)

**Table 2.** Metrics of peri-movement SDF. (mean ± SEM, ** p<0.01 between VLp and VLa, † p<0.05, †† p<0.01 between motor thalamic nuclei and M1, Wilcoxon rank test)

|  |  | VLp | VL<sub>a</sub> | M1 |
| --- | --- | --- | --- | --- |
| Increase only | Latency (ms) | 112.5 ± 9.3 ** | 82.4 ± 7.0 † | 104.3 ± 6.6 |
| | Magnitude (Hz) | 16.7 ± 1.4 †† | 13.2 ± 0.8 †† | 34.2 ± 2.2 |
| | Slope (Hz/ms) | 0.16 ± 0.01 †† | 0.15 ± 0.01 †† | 0.35 ± 0.02 |
| | Half width (ms) | 248.2 ± 26.1 †† | 257.0 ± 22.0 †† | 199.5 ± 18.1 |
| Decrease only | Latency (ms) | 58.6 ± 23.7 | 71.9 ± 8.6 | 87.6 ± 13.7 |
| | Magnitude (Hz) | 9.1 ± 1.2 † | 9.4 ± 0.5 †† | 12.8 ± 1.0 |
| | Slope (Hz/ms) | 0.13 ± 0.02 † | 0.11 ± 0.01 †† | 0.18 ± 0.02 |
| | Half width (ms) | 342.8 ± 98.4 | 472.8 ± 54.9 | 333.5 ± 64.6 |

For increase-type responses, only one measure differed significantly between VLp and VLa populations: responses began earlier relative to movement onset in VLp than in VLa (mean 30 msec earlier; Wilcoxon rank test; p<0.01; Table 2). Increase-type responses in VLp and in M1 began at similar pre-movement onset times. The magnitudes and durations of increase-type responses were statistically indistinguishable between VLp and VLa (Wilcoxon rank test; ps≥0.1), while those in M1 were markedly larger and shorter duration (Wilcoxon rank test; ps<0.01; Table 2, *Magnitude, Slope and Half-width*).

For decrease-type responses, onset latencies did not differ significantly between the two thalamic nuclei or relative to M1. Similar to what was seen for increase-type responses, response magnitudes were indistinguishable between VLp and VLa while those in M1 were larger (Wilcoxon rank test; all p<0.05; Table 2, *Magnitude* and *Slope*). Finally, the duration of decrease-type responses was prolonged in VLa relative to those in M1 neurons (Wilcoxon rank test; p<0.01), while in VLp the duration of decreases was intermediate between VLa and M1 (Wilcoxon rank test; p=0.16).

To summarize, we found two notable exceptions to our general prediction that peri-movement responses do not differ between VLp and VLa (Nambu et al., 1988; Anderson and Turner, 1991; van Donkelaar et al., 1999). First, increase-type responses began earlier in VLp than they did in VLa. That result is consistent with the idea that movement-related signals in the cerebello-thalamocortical circuit have an early influence on the timing of movement initiation (Ivanusic et al., 2005; Nashef et al., 2019; Dacre et al., 2021), whereas signals in the BG-thalamocortical circuit do not (Goldberg and Fee, 2012; Schwab et al., 2020). We also found that decrease-types responses were more common in VLa than in VLp. That difference might be a product of the strong inhibitory input that VLa receives from GPi. For the most part, however, peri-movement responses in VLp and VLa were statistically indistinguishable, contrary to what might be predicted from the categorically-different sub-cortical inputs that the two nuclei receive. We next tested whether the activity of VLp and VLa might be more readily differentiated by the dimensions of task performance encoded by their neuronal activity or by the timing of that neural encoding.

### 3.5 Time-resolved regression identified differences in encoding

We used time-resolved linear regression analyses to determine if peri-movement activity differed between VLp, VLa, and M1 with respect to the dimensions of task performance encoded or the timing of that encoding. Between-trial variations in a unit’s peri-movement firing rate were regressed against the following trial-by-trial measures of task performance: 1) duration of the start position hold period (prior to presentation of the go-cue), 2) reaction time, 3) mean movement speed, 4) reach direction, and 5) elapsed time from the beginning of the recording session (“session time”; see *Methods*). The collinearity between predictors was acceptably low (Belsey’s collinearity: 3.39, 4.18 and 2.49 for predictors from VLp, VLa and M1 recording sessions, respectively (Belsley, 1991). Trial-by-trial time-evolving estimates of firing rate were regressed against those five predictors. All units were included in this analysis regardless of the response type observed.

The regressions identified relationships between one or more predictors in large proportions of all neural populations (63%, 53% and 74% of units in VLp, VLa and M1, chi-square test; *χ^2^*(2, n=448) = 17.33, p<0.01). Typical regression results are shown for a single VLp unit in Fig. 3A. During the start position hold period, prior to go-cue presentation (Fig. 3A; median shown by *vertical dashed line* and IQR by *gray* shading), the full model accounted for only a small fraction of the variance in firing rate (5% approx.; R^2^ for full model, *black trace* Fig. 3A top), essentially all of which was attributable to a significant relationship between firing rate and the time elapsed in the recording session (semi-partial R^2^, “pR^2^”, for session time, *red trace* Fig. 3A top). Small but consistently negative beta values for that relationship between firing rate and session time (*red trace* Fig. 3A bottom) indicated that firing rates during this period declined slowly across the whole recording session. Other regressors accounted for negligible fractions of firing rate variance during the delay period. Immediately following the median time of go-cue presentation (240 +/- 32 ms prior to movement onset) the full model R^2^ increased sharply and reached a peak of 26% of variance at 154 ms prior to movement onset. That increase was attributable to a strengthening of the relationship between firing rate and session time and the appearance of a strong relationship to movement direction (pR^2^ for direction; *blue trace* Fig. 3A top). A second, long-lasting elevation in R^2^, which began 50 ms before movement onset, was composed of successive peaks in pR^2^ for movement direction, movement speed (*yellow trace*), delay period (*gray trace*) and then a third large peak for direction (*blue*). Regression results for RT are not shown in Fig. 3A because the pR^2^ for RT was zero in this unit.

**Figure 3.**
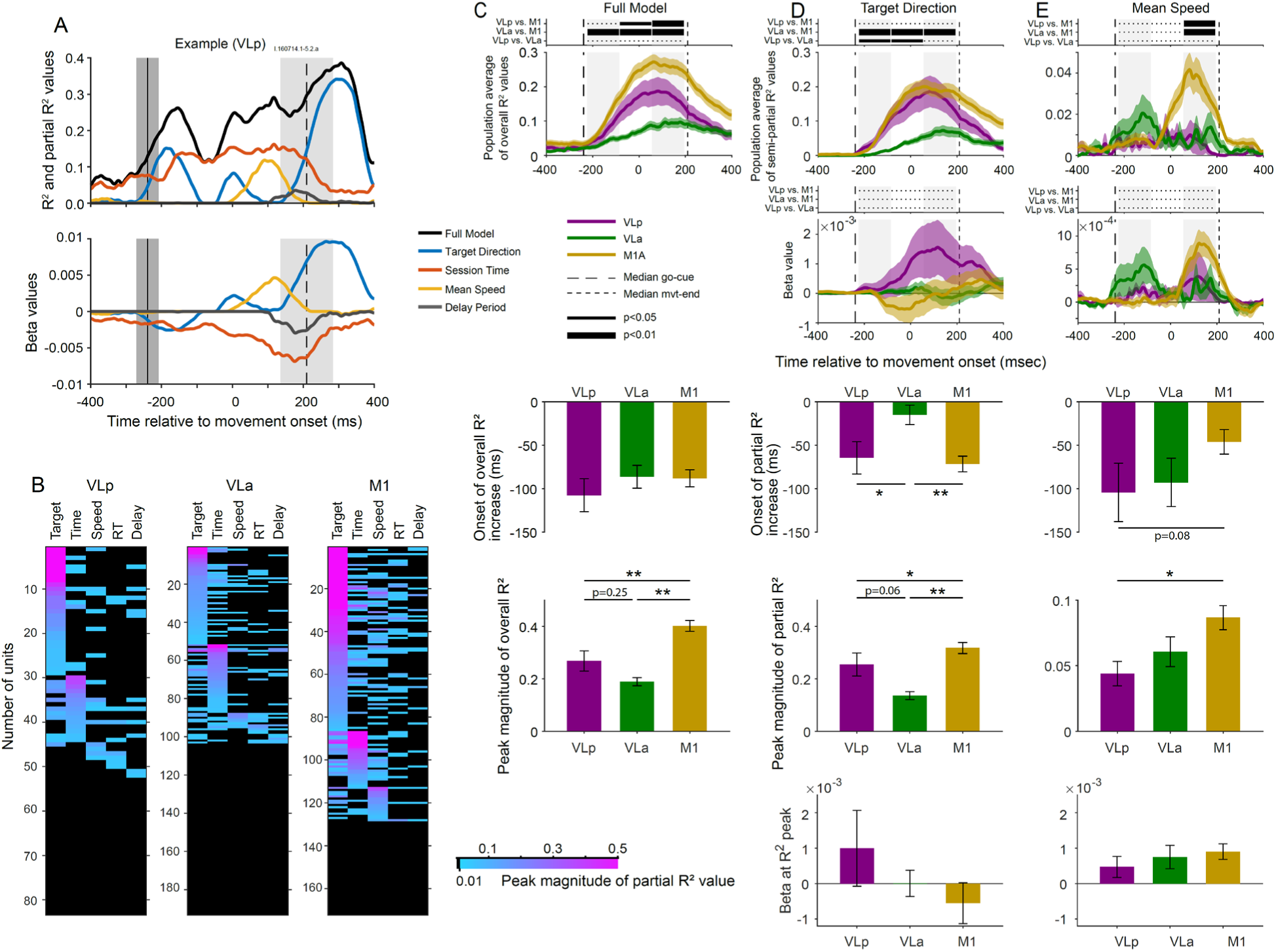
**(A)** Exemplar regression results for activity of a single VLp neurons influenced by multiple predictors. Top: R² values. Bottom: beta values. Black: full model; orange: mean speed; magenta: target direction; blue: delay period; green: session time. Vertical black dashed lines and gray rectangles mark the median and interquartile range (IQR) of go-cue onset times (left) and movement end times (right), aligned to movement onset. **(B)** Peak semi-partial R² values for all single units, sorted according to the most effective predictor and peak magnitude (larger semi-partial R² values at the top within each predictor category). **(C)** Population averages of R² values for VLp (magenta), VLa (green), and M1 (yellow). *Shaded areas*: ±SEM. **(D–E)** Population averages of semi-partial R² values (top) and beta values (bottom) associated with target direction (D) and mean speed (E). Vertical black dashed lines indicate the median times of go-cue onset (left) and movement end (right). Shaded areas represent ±SEM. Horizontal lines above each panel (C-E) indicate Wilcoxon rank test results for early, middle, and late epochs of the response interval (thin black line: p<0.05; thick black line: p<0.01; Wilcoxon test with p-values corrected for multiple comparisons using false discovery rate; horizontal dotted line: not significant).

We analyzed the number and combination of predictors that showed significant increases in pR^2^ values during the peri-movement test period (“test period”; see *Methods*). The magnitudes of peak pR^2^ values of all predictors for all units are summarized in heat maps (Fig. 3B), where each row represents the pR^2^ coefficients for a single unit for each of the five predictors (columns). Target direction was the most common predictor for all neural populations (Table 1 *Behavior encoding units*). Similar fractions of neurons encoded target direction in VLp and VLa populations (47% and 40% respectively; chi-square test; *χ^2^*(1, n=276) = 1.20, p=0.27) although those fractions were smaller than the fraction of M1 neurons found to encode direction (66%; chi-square test; *χ^2^*(2, n=448) = 26.11, p<0.01). Progressively smaller fractions of neurons in each population encoded session time and movement speed. Only a small number of neurons showed significant effects for RT and delay predictors. We found no consistent pattern to the way encodings of multiple predictors were distributed across populations. Therefore, from here-on we will focus primarily on results for the full model and the target direction and movement speed regressors.

To compare the timing and strength of task encoding, we constructed time-resolved population averages of overall R^2^ values for the three neural populations. Those curves began to rise at approximately 200 ms before movement onset for all neural populations (Fig. 3C top).

Encoding was stronger in VLp than in VLa throughout the reaction time and move execution periods. Interestingly, task encoding was stronger in the M1 population than in either of the thalamic areas (Fig. 3C top). The times of onset of significant encoding averaged across individual neurons confirmed that the onset of encoding did not differ between populations (Wilcoxon rank test; between all pairs; ps>0.23; Fig. 3C middle). Neuron-by-neuron measures of the peak magnitude of overall R^2^ also confirmed the observation that overall encoding of task parameters was weaker in both VLp and VLa as compared with the M1 population (Wilcoxon rank test; ps<0.01; Fig. 3C bottom). Interestingly however, when we extracted the magnitudes of peak R^2^ neuron-by-neuron (irrespective of the timing of the peak), we found no significant difference between VLp and VLa populations (Wilcoxon rank test; p=0.24), contrary to what the population average curves in Fig. 3C predict. We revisit this disparity in the next section.

Next, to identify potential differences between brain areas in their relationship to the encoding of target direction, we computed pR^2^ and beta values separately for each neural population and compared the onsets and peak magnitudes of significant encoding between VLp, VLa and M1 populations (Fig. 3D).

### 3.6 Encoding of direction appears earlier and is stronger in VLp than in VLa

For both VLp and M1 populations, the population averages of the pR² values for target direction were similar to each other with respect to time course and magnitude of change. The curves of pR² values began to rise approximately 200 ms before movement onset (Fig. 3D top, *magenta* and *yellow*). In contrast, for VLa the appearance and growth of direction encoding was more gradual, lagging well behind that of VLp and M1 (Figs. 3D top, green). Comparison of the mean times of onset of direction encoding across individual neurons confirmed that in VLa the appearance of direction encoding was delayed relative to that in VLp or M1 (Wilcoxon rank test; p < 0.05 for VLp, p < 0.01 for M1). The strength of direction encoding was also similar for VLp and M1.

When comparing overall R² values and semi-partial R² values for target direction to evaluate the contribution of target direction encoding to overall encoding, we found that VLp and VLa showed similar magnitudes of change between these metrics (Fig. 3D top, magenta and green). In contrast, in M1, the changes in semi-partial R² values were smaller than those in overall R² values, suggesting that other predictors had a greater influence on overall R² in M1 compared to VLp and VLa (Fig. 3D top, yellow). The close alignment of overall and semi-partial R² magnitudes in VLp and VLa suggests that target direction encoding was the primary contributor to the overall R² in these regions. Neuron-by-neuron measurements of peak semi-partial R² magnitudes revealed significant differences between M1 and VLp and VLa (Fig. 3D second from bottom). Specifically, the semi-partial R² magnitudes in both VLp and VLa populations were significantly smaller than in the M1 population (Wilcoxon rank test; p < 0.05 for VLp, p < 0.01 for VLa). However, despite the difference in population averages of semi-partial R² values between VLp and VLa, this difference was not statistically significant at the individual neuron level (Wilcoxon rank test; p = 0.06).

### 3.7 Encoding of mean speed begins early in VLa

The population averages of the semi-partial R² values for mean speed, derived from the speed-encoding neuron population, were different from the population averages of overall R² values in VLp, VLa and M1. The curves of semi-partial R² values began to rise approximately 200 ms before movement onset, but the mean magnitude was smaller in VLp compared to VLa and M1 (Fig. 3E top, magenta). In VLa, the semi-partial R² value for mean speed began increasing immediately after go-cue presentation, with strong encoding observed in the reaction time (Fig. 3E top, green). In contrast, mean speed encoding in M1 was stronger during the reach phase (Fig. 3E top, yellow). Quantitative analyses of the onset of significant encoding and the magnitude of semi-partial R^2^ value in individual neurons aligned with trends observed in the population averages of semi-partial R^2^ and beta values (Fig. 3E middle to bottom).

### 3.8 Clustering analysis identified two types of target direction encoding in VLp

In the population-averaged curves of both overall and semi-partial R² values for target direction, we observed clearly distinct encoding patterns between VLp and VLa populations (Figs. 3C top and 2D top). However, neuron-by-neuron analysis of peak encoding magnitudes did not show statistically significant differences (Figs. 4A). We hypothesized that this discrepancy might arise from large variability in encoding magnitudes within the VLp population. To test this hypothesis, we compared the distributions of peak magnitudes for semi-partial R² values of target direction across the three populations (Fig. 4A). The cumulative distribution functions (CDFs) for semi-partial R² values in VLa and M1 populations displayed smooth, logarithmic curves (Fig. 4A, green and yellow). In contrast, the CDFs in the VLp population showed a plateau in the middle, suggesting that the magnitude distribution in VLp may arise from multiple underlying sources (Fig. 4A, magenta).

**Figure 4.**
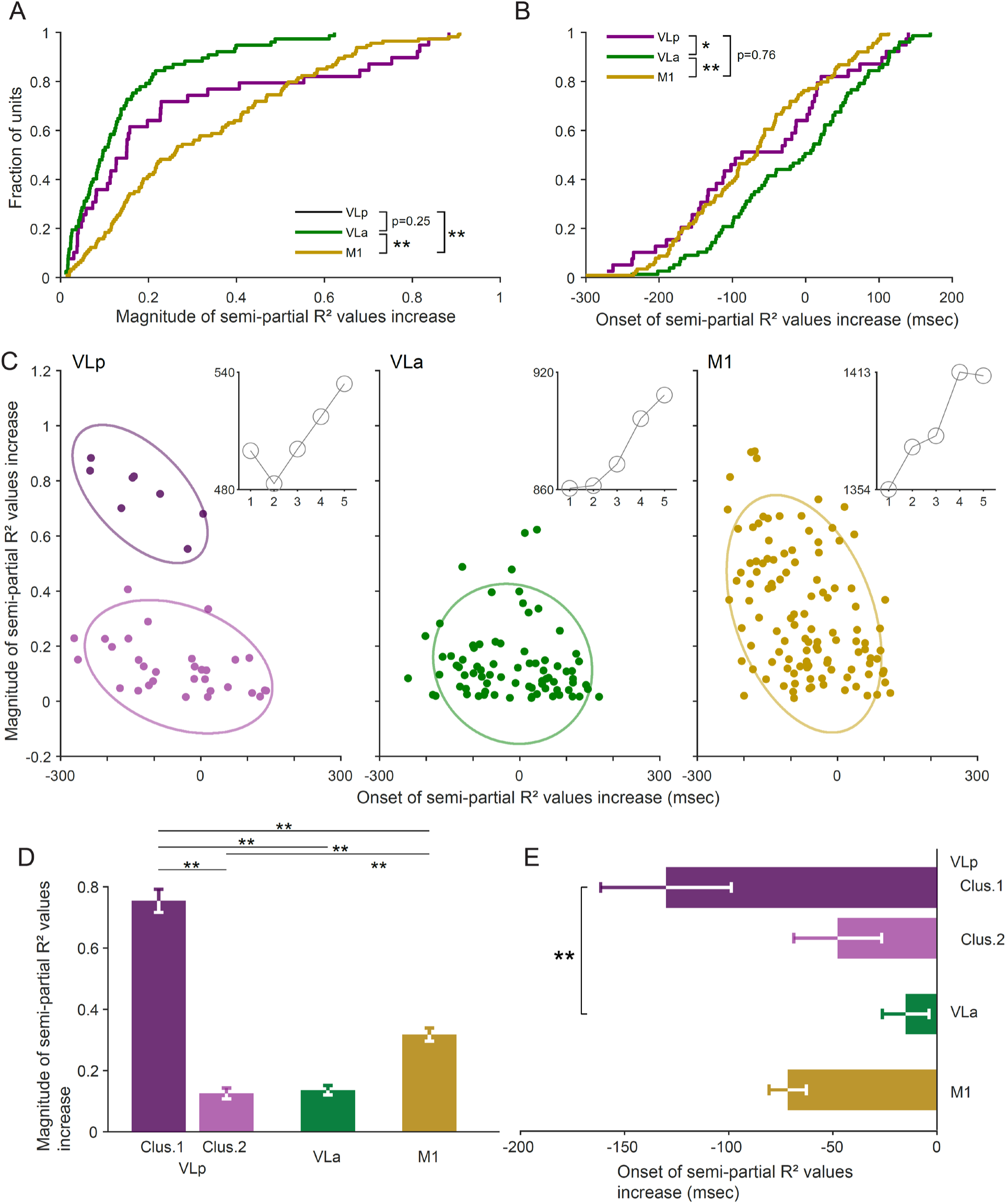
**(A)** Cumulative distributions of the peak magnitudes of semi-partial R² values associated with target direction in VLp (magenta), VLa (green), and M1 (yellow). Peak magnitudes in M1 were significantly larger than those in VLp and VLa (** p < 0.01, Wilcoxon rank test), whereas no significant difference was observed between VLp and VLa. **(B)** Cumulative distributions of the onsets of change in semi-partial R² values associated with target direction in VLp (magenta), VLa (green), and M1 (yellow). Onsets in VLa were significantly delayed relative to VLp and M1 (* p < 0.05, ** p < 0.01, Wilcoxon rank test), whereas no significant difference was found between VLp and M1. **(C)** Scatter plots of the onset of change in semi-partial R² values associated with target direction (x-axis) versus the magnitude of change (y-axis) for VLp (left), VLa (middle), and M1 (right). Insets: Bayesian Information Criterion (BIC) values from clustering analyses. X-axis indicates the number of clusters used. **(D and E)** Mean magnitude (D) and mean onset (E) of the significant increase in semi-partial R² values associated with target direction (** p < 0.01, Wilcoxon rank test)

The presence of multiple subpopulations with varying peak magnitudes of semi-partial R² in VLp may help explain the high p-values observed in our statistical tests. We focused on target direction encoding because most of the overall R² values in VLp and VLa arise from the semi-partial R² values associated with target direction (Figs. 3C top and 3D top, magenta and green). In addition to the cumulative distribution function (CDF) of semi-partial R² magnitude, we also observed a plateau in the CDF of encoding onset in VLp (Fig. 4B, magenta).

To further analyze the distribution of peak semi-partial R² magnitudes within the VLp population, we created a scatter plot showing the relationship between peak semi-partial R² magnitude and encoding onset (Fig. 4C left). This plot suggested two clusters within VLp: one with high peak semi-partial R² values and early encoding onset, and another with low peak magnitudes and a broad range of encoding onsets. To test this, we fitted Gaussian mixture models to the distributions of both peak magnitudes and encoding onsets, testing models with 1 to 5 clusters. The optimal number of clusters was determined by comparing the Bayesian Information Criterion (BIC), with a smaller BIC indicating a better fit. In VLp, the two-cluster model yielded the lowest BIC (Fig. 4C left inset). In the scatter plot, markers are color-coded to indicate cluster membership (Fig. 4C). One VLp cluster demonstrated early and strong target direction encoding (Cluster 1: dark magenta markers), while the other displayed a wide range of onset times and weaker encoding (Cluster 2: light magenta markers). Similar clustering analyses for VLa and M1 populations suggested single, unified clusters in both regions (Fig. 4C middle and right).

Finally, we analyzed the metrics of clusters 1 and 2 in the VLp population and compared them with those in VLa and M1. The magnitude of encoding in VLp cluster 1 neurons was significantly greater than those in all other populations (Wilcoxon rank test; all p < 0.01; Fig. 4D). The onset of encoding in VLp cluster 1 neurons was also significantly earlier than in VLa (Wilcoxon rank test; p < 0.05; Fig. 4E). Conversely, the magnitude of encoding in VLp cluster 2 neurons did not differ significantly from that in VLa but was significantly smaller than in M1 (Wilcoxon rank test; p = 0.89 for VLa and p < 0.01 for M1; Fig. 4D). Similarly, the encoding onset of VLp cluster 2 neurons was not significantly different from that in either VLa or M1 (Wilcoxon rank test; p = 0.15 for VLa and p = 0.27 for M1; Fig. 4E).

We observed similar plateau in the CDF of magnitude of semi-partial R^2^ values for mean speed in VLa (Fig. 4B, magenta). However, clustering analyses for mean speed encoding did not reveal a multi-cluster structure in any of the regions analyzed.

Although VLp and VLa did not significantly differ in peri-movement SDF, linear regression analysis indicated that VLp neurons tended to encode target direction earlier and more strongly than VLa neurons. Our clustering analysis using Gaussian model fitting showed that a two-cluster model produced the smallest BIC for target direction-encoding VLp neurons. This result suggests that target direction-encoding neurons in VLp can be divided into two groups: one group that begins encoding target direction during the reaction time with strong encoding, and another group that encodes target direction weakly, with encoding onset distributed throughout the response time. This clustering pattern was not observed in VLa or M1 and appears to be a unique property of VLp.

## 4 Discussion

We compared neuronal activity in VLp and VLa during a choice reaction-time reaching task. Consistent with previous reports, the trial-averaged peri-movement activity in the two areas was broadly similar. However, we found that the proportion of decrease-only responses was significantly larger in VLa than in VLp, and that VLp neurons encoded movement direction more strongly than VLa neurons. In addition, we identified a subpopulation of VLp neurons that exhibited particularly robust direction encoding during the reaction period. Together, these findings replicate the well-established similarity in peri-movement responses while revealing, for the first time, a functional distinction between VLp and VLa. This VLp subpopulation may play a key role in motor-control mechanisms mediated through the cortico-cerebellar–VLp loop.

### Strong inhibitory input in VLa

Although peri-movement activity patterns in VLp and VLa were previously reported to be similar (Anderson and Turner, 1991; van Donkelaar et al., 1999), our analysis revealed several indications of strong inhibitory input in VLa. Specifically, the proportion of neurons exhibiting a monophasic decrease in firing rate around movement was significantly higher in VLa compared to VLp (Fig. 2D). Quantitative analysis of the spike density function (SDF) revealed a prolonged decrease in VLa, although the duration was not significantly longer than that observed in VLp (Table 2). These findings suggest that VLa neurons receive stronger inhibitory input than VLp neurons.

The globus pallidus internus (GPi) is the most likely source of this inhibitory input to VLa. This hypothesis is supported by the unique giant synaptic structures formed by inhibitory projections from the basal ganglia output nuclei to the motor thalamus (Ilinsky et al., 1997; Bodor et al., 2008; Rovo et al., 2012) and by studies demonstrating inhibitory effects of GPi electrical stimulation on thalamic activity (Uno et al., 1970; Nambu et al., 1991; Ando et al., 1995; Kim et al., 2017). Other GABAergic sources, such as the reticular thalamic nucleus (nRT) and local interneurons, may also contribute to the stronger inhibitory input in VLa. However, these GABAergic projections are known to target VLp as well, and previous studies suggest that nRT and interneuron inputs may actually be stronger in VLp than in VLa. For instance, axons from motor nRT neurons predominantly form synapses on the distal dendrites of thalamocortical neurons in VLa, whereas in VLp, nRT axons synapse directly on the soma and proximal dendrites (Ilinsky et al., 1999). Additionally, cortico-thalamic projections from M1 form synapses on the dendrites of interneurons in VLa but form presynaptic terminals of interneurons in VLp (Galvan et al., 2016).

Thus, given the relatively weaker inhibitory contributions from nRT and local interneurons in VLa, the greater prevalence of decrease-type peri-movement responses is most likely a product of the strong inhibitory input to VLa from the GPi.

### Small fraction of VLp neuron encode the target direction strongly

Previous studies reported similar activity patterns in VLp and VLa during movement. Those studies, however, did not test for differences between nuclei in the neural encoding of task parameters. To address this, we performed time-resolved linear regression analyses using behavioral parameters. Among these, target direction emerged as the most influential predictor across all examined target areas (Fig. 3B). This result aligns with prior findings of target direction encoding in the regions upstream of VLp and VLa [e.g., the deep cerebellar nuclei (Fu et al., 1997) and GPi (Turner and Anderson, 1997)].

The distribution of target direction encoding differed, however, between VLp and VLa (Fig. 4C). Clustering analysis revealed that the best-fitting model for VLp target direction encoding units was a two-cluster model (based on the lowest BIC), whereas the same analysis for VLa units supported a single-cluster model. Using the clustering results, we recalculated the metrics of target direction encoding and found that the subgroup of VLp neurons with smaller partial R² values for target direction were indistinguishable from VLa neurons in their encoding metrics (Fig. 4D-E).

These findings suggest that only a subset of VLp neurons receives strong target-direction-related input from their upstream brain areas.

### Potential encoding of behavioral parameter by a few impactful thalamic neurons

Similar to the distribution of partial R² values for target direction, we observed an irregular CDF for partial R² values associated with mean speed in VLa (Supplemental Fig. S2B, top). The encoding of mean speed in the VLa aligns with previous studies that demonstrated the effect of transient GPi inactivation on movement vigor ((Turner and Desmurget, 2010) and references therein). Additionally, the observation that only a small fraction of VLa neurons strongly encode mean speed is consistent with a prior report of peak speed encoding in a subset of GP neurons (Turner and Anderson, 2005). However, our clustering analysis did not identify multi-cluster models as the best fit for this dataset. This result of clustering analysis may be due to the small number of VLa neurons that strongly encoded mean speed, despite a total sample size of 193 VLa neurons, which is not insubstantial.

One potential implication of these findings is that both strong target direction encoding in VLp and strong speed encoding in VLa are represented in small subpopulations of neurons within these recipient VL nuclei. This hypothesis is supported by histological evidence showing that axons from both the dentate nucleus and GPi form focal terminal fields in their target regions (Arecchi-Bouchhioua et al., 1996; Ilinsky et al., 1997; Mason et al., 2000). These studies suggest that individual axons may selectively influence small clusters of neurons.

This model could also explain the consistently small differences between VLp and VLa when compared at the whole population level. Future research using high-density probes capable of capturing neural activity from neurons in these small clusters may provide a more definitive understanding of these patterns.

## Conclusion

The anatomical separation of Cb- and BG-recipient thalamic neurons suggests that the distinct roles of Cb- and BG-thalamocortical circuits in motor control should be evident when comparing the movement-related activities of neurons in VLp and VLa. Surprisingly, however, previous studies reported similar peri-movement activity in these nuclei during behavioral tasks (Anderson and Turner, 1991; van Donkelaar et al., 1999). Our analysis revealed that the fraction of neurons showing a decrease-only response was significantly larger in VLa than in VLp. While quantitative analyses of peri-movement activity indicated a trend toward stronger peri-movement increase in firing rate in VLp and stronger peri-movement decrease in firing rate in VLa, these differences were not statistically significant.

In contrast, our regression and clustering analyses highlighted differences in the encoding of the behavioral parameter associated with target direction between VLp and VLa. Specifically, we identified two groups of VLp neurons with differing degrees of target direction encoding. In contrast, the same clustering analysis suggested a single-cluster distribution for VLa neurons.

One group of VLp neurons exhibited strong early target direction encoding during the reaction time, while the other group’s encoding was indistinguishable from the target direction encoding observed in VLa.

Future studies employing higher spatial resolution recording techniques and pharmacological inactivation of upstream brain areas will be critical for further elucidation of the functional differences between VLp and VLa within the motor control system.

## 6 Author Contributions

RST conceived and designed research; DK, AJZ, and RST performed experiments; DK and KC analyzed data; DK, KC, TP, and RST interpreted results of experiments; DK prepared figures; DK, and RST drafted manuscript; DK, AJZ, KC, TP, and RST edited and revised manuscript and approved final version of manuscript.

## Acknowledgment

We thank Lisa Nieman-Vento and Cherie Lee Cornmesser for their contributions to animal care.

## 8 Data Availability

Both electrophysiological data and behavioral data are available on the DANDI Archive at DANDI:000947/0.240510.2211. All MatLab code and the processed source data sufficient to reproduce all figures and tables are in preparation and will be available on Zenodo (<u>TO</u> BE FILLED).

## 9 Key Resource Table

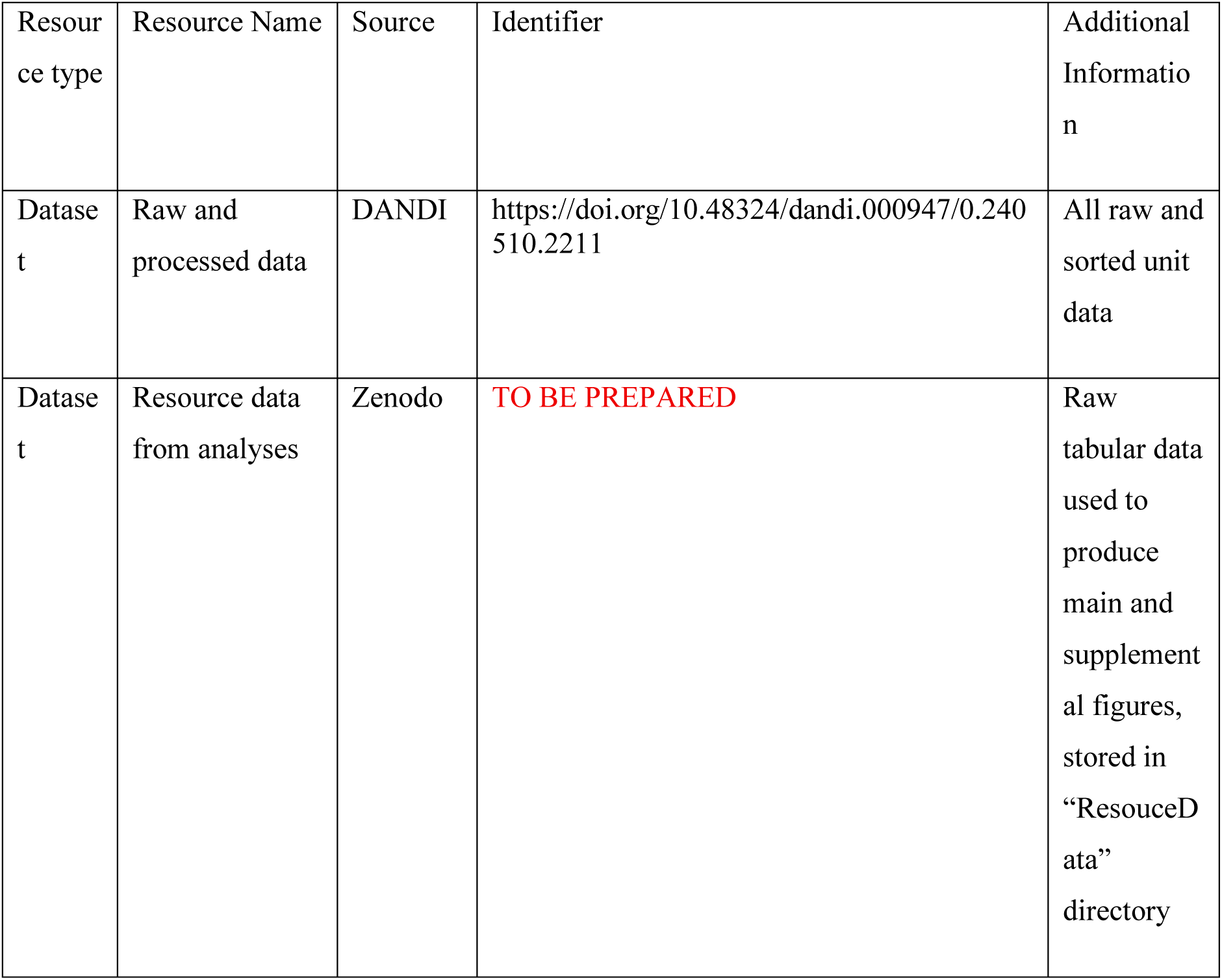
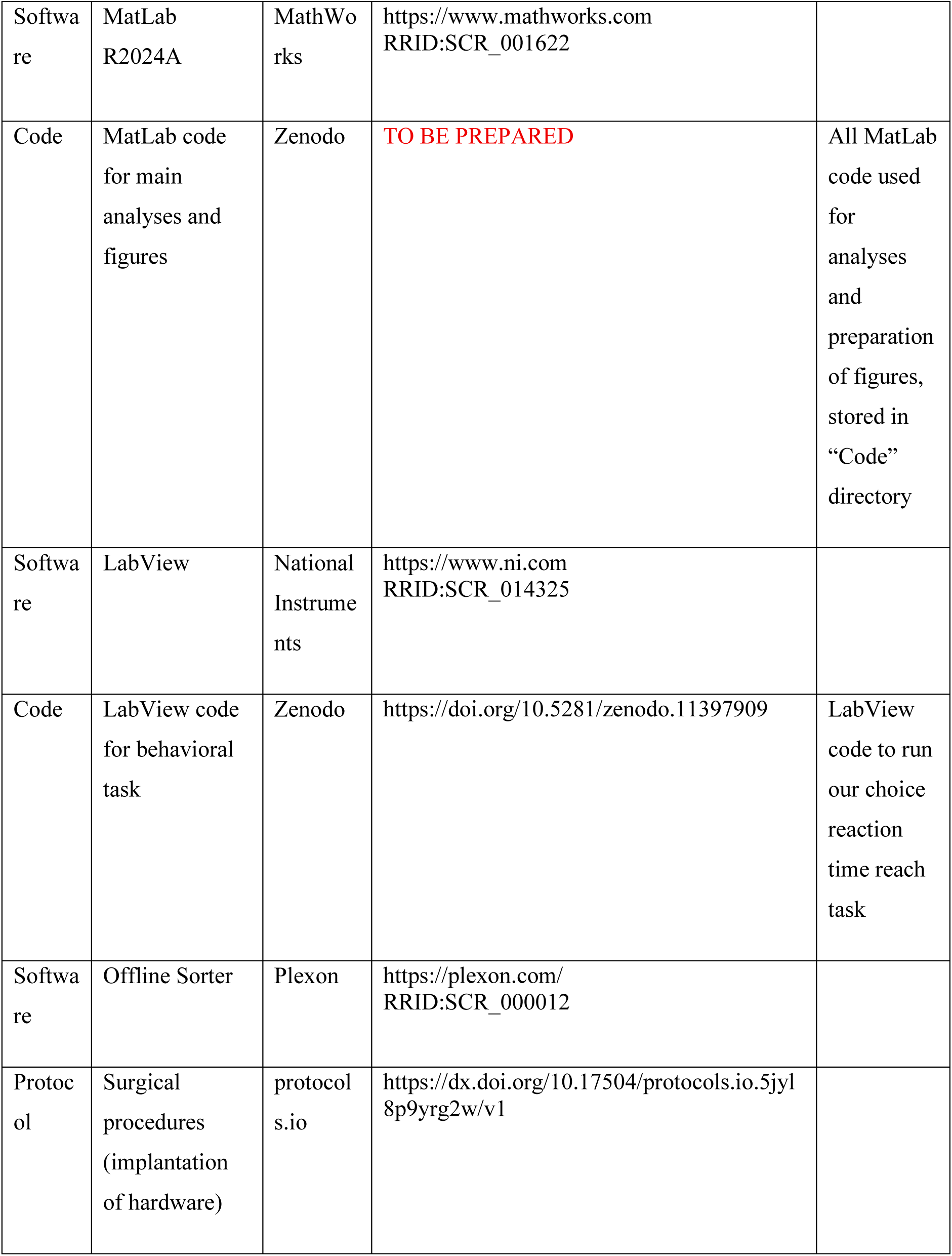
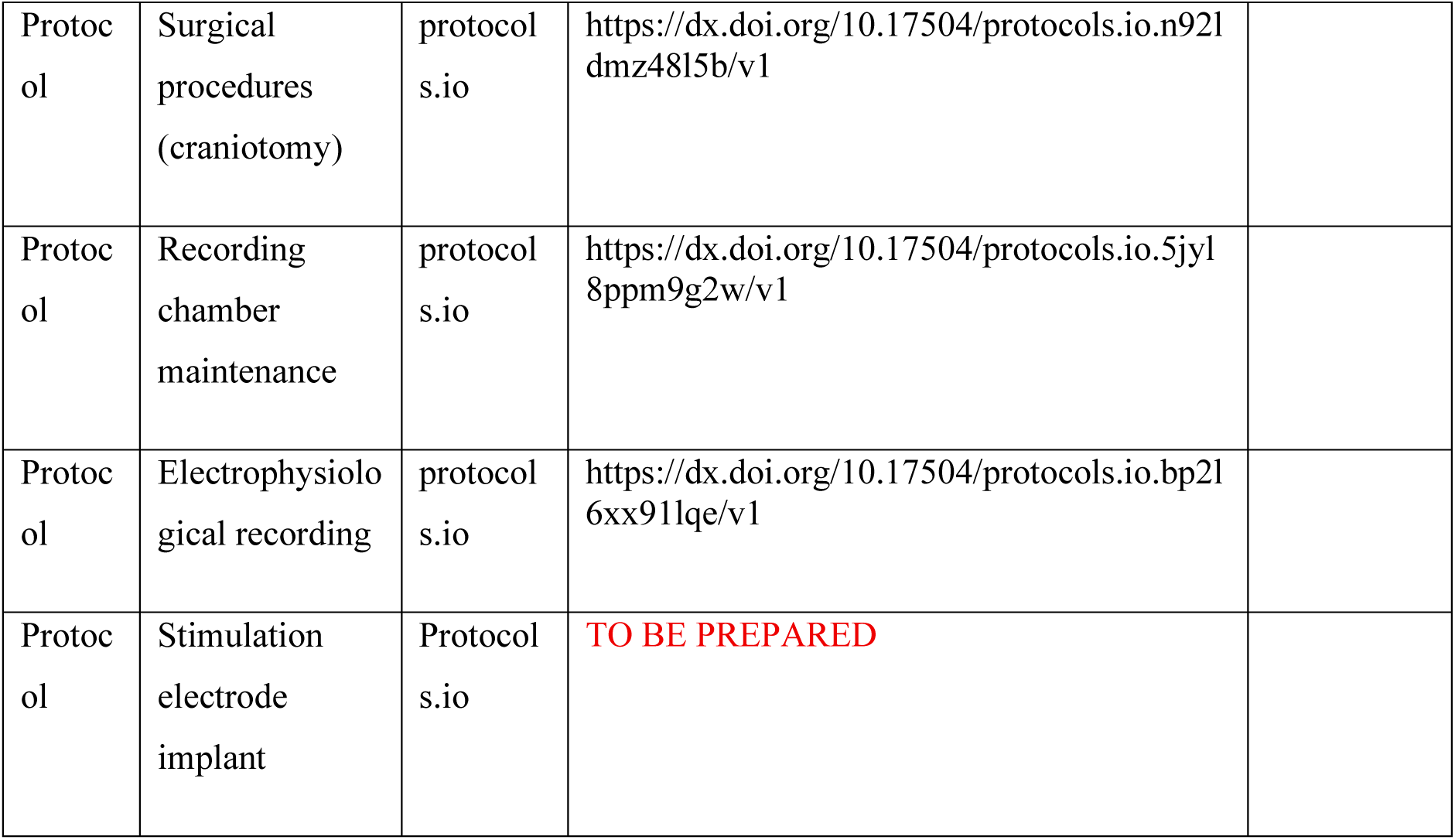

10 Key Resource Table
| Resource type | Resource Name | Source | Identifier | Additional Information |
| --- | --- | --- | --- | --- |
| Dataset | Raw and processed data | DANDI | https://doi.org/10.48324/dandi.000947/0.240510.2211 | All raw and sorted unit data |
| Dataset | Resource data from analyses | Zenodo | **TO BE PREPARED** | Raw tabular data used to produce main and supplemental figures, stored in “ResourceData” directory |
| Software | MatLab R2024A | MathWorks | https://www.mathworks.com<br>RRID:SCR_001622 |  |
| Code | MatLab code for main analyses and figures | Zenodo | TO BE PREPARED | All MatLab code used for analyses and preparation of figures, stored in “Code” directory |
| Software | LabView | National Instruments | https://www.ni.com<br>RRID:SCR_014325 |  |
| Code | LabView code for behavioral task | Zenodo | https://doi.org/10.5281/zenodo.11397909 | LabView code to run our choice reaction time reach task |
| Software | Offline Sorter | Plexon | https://plexon.com/<br>RRID:SCR_000012 |  |
| Protocol | Surgical procedures (implantation of hardware) | protocols.io | https://dx.doi.org/10.17504/protocols.io.5jyl8p9yrg2w/v1 |  |
| Protocol | Surgical procedures (craniotomy) | protocol s.io | https://dx.doi.org/10.17504/protocols.io.n921dmz48l5b/v1 |  |
| Protocol | Recording chamber maintenance | protocol s.io | https://dx.doi.org/10.17504/protocols.io.5jyl8ppm9g2w/v1 |  |
| Protocol | Electrophysiological recording | protocol s.io | https://dx.doi.org/10.17504/protocols.io.bp2l6xx9l1qe/v1 |  |
| Protocol | Stimulation electrode implant | Protocol s.io | TO BE PREPARED |  |

---

**Supplementary Figure 1.**
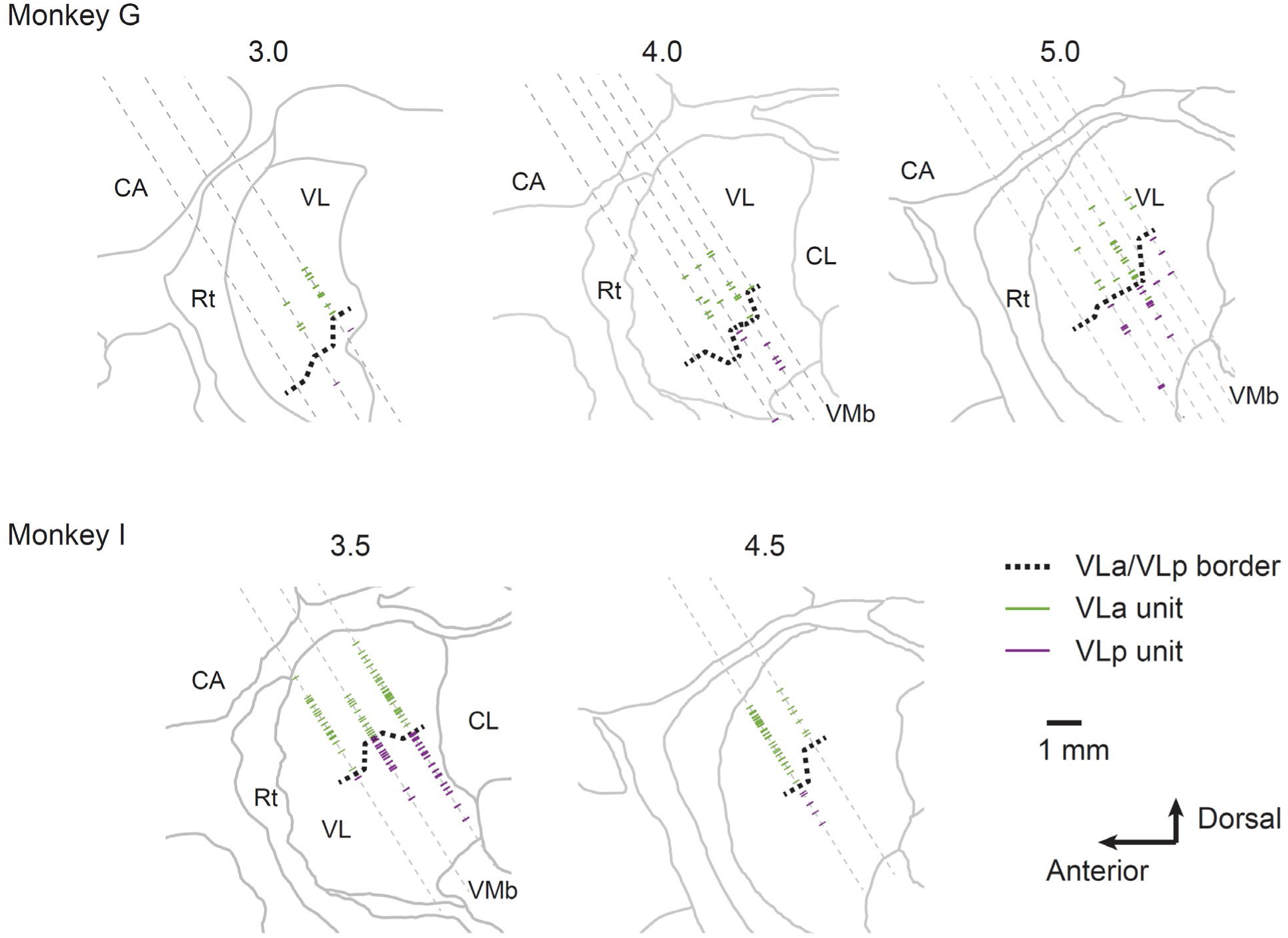
Reconstructed locations of all VLp (magenta) and VLa (green) single units included in the database. The VLp–VLa boundary (black dashed lines) was defined by responses to SCP and GPi stimulation during border-mapping sessions prior to data collection. Locations are projected onto parasagittal sections at 1 mm intervals for animals G and I separately. The boundary of thalamus was taken from a standard atlas and aligned with the mapping results from each animal.

**Supplementary Figure 2.**
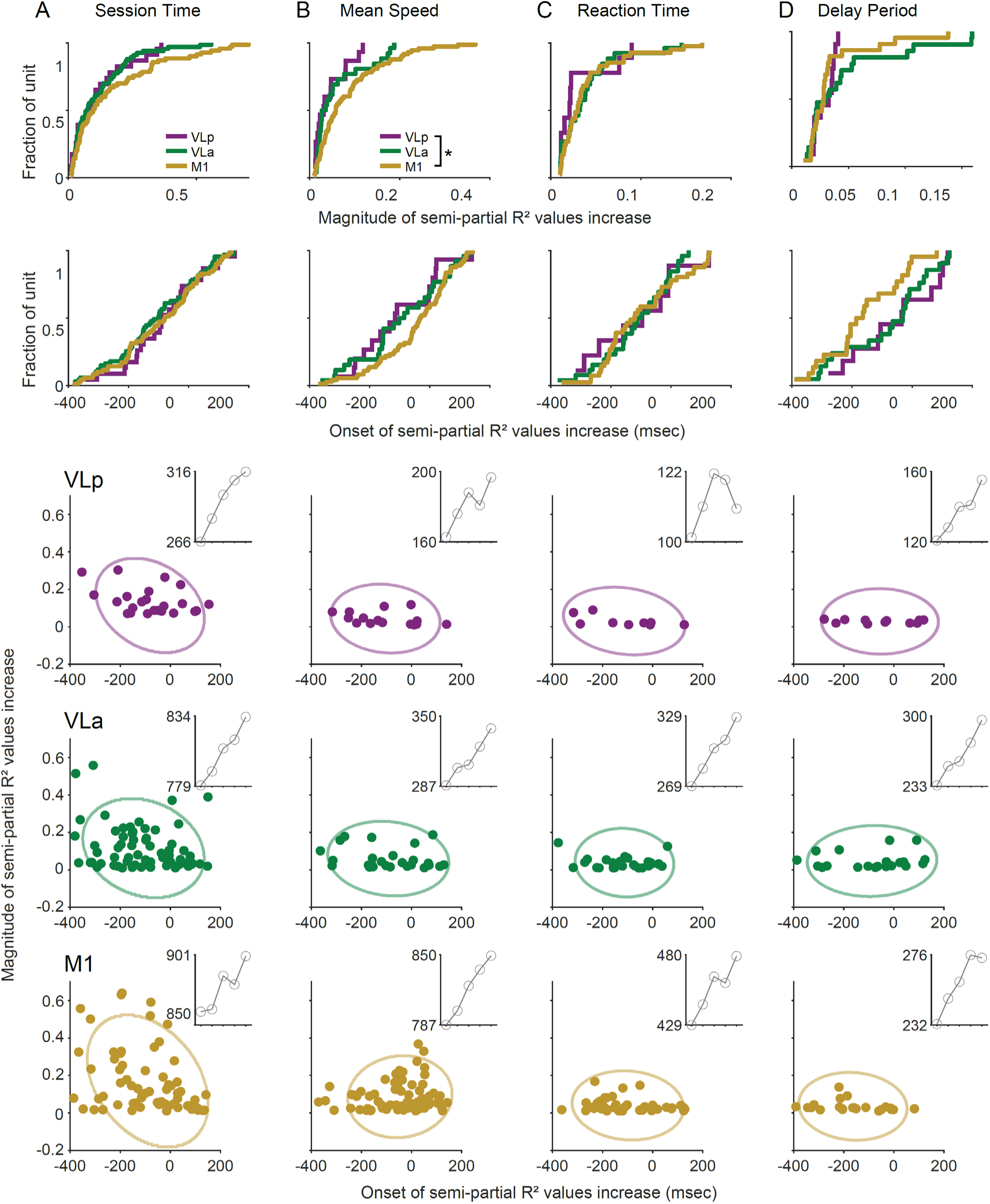
**(A–D)** Top and second rows: cumulative distributions of the peak magnitudes (top row) and onsets (second row) of partial R² values associated with session time, mean speed, reaction time, and delay period (from left to right). Partial R² values associated with mean speed in VLp were significantly smaller than those in M1 (* p < 0.05, Wilcoxon rank test). Third, fourth, and bottom rows: scatter plots of the onset of change in partial R² values associated with predictors (x-axis, session time, mean speed, reaction time, and delay period, from left to right) versus the magnitude of change (y-axis) for VLp (third row), VLa (fourth row), and M1 (bottom row). Insets: BIC values from clustering analyses. X-axis indicates the number of clusters used.

